# An ancient role for Collier/Olf/Ebf (COE)-type transcription factors in axial motor neuron development

**DOI:** 10.1101/454926

**Authors:** Catarina Catela, Edgar Correa, Jihad Aburas, Laura Croci, G. Giacomo Consalez, Paschalis Kratsios

## Abstract

**Background:** Mammalian motor circuits display remarkable cellular diversity with hundreds of motor neuron (MN) subtypes innervating hundreds of different muscles. Extensive research on limb muscle-innervating MNs has begun to elucidate the genetic programs that control animal locomotion. In striking contrast, the molecular mechanisms underlying the development of axial muscle-innervating MNs, which control breathing and spinal alignment, are poorly studied.

**Methods:** Our previous studies indicated that the function of the Collier/Olf/Ebf (COE) family of transcription factors (TFs) in axial MN development may be conserved from nematodes to simple chordates. Here, we examine the expression pattern of all four mouse COE family members (mEbf1-mEbf4) in spinal MNs and employ genetic approaches in both nematodes and mice to investigate their function in axial MN development.

**Results:** We report that mEbf1 and mEbf2 are expressed in distinct MN clusters (termed “columns”) that innervate different axial muscles. Mouse Ebf1 is expressed in MNs of the hypaxial motor column (HMC), which is necessary for breathing, while mEbf2 is expressed in MNs of the medial motor column (MMC) that control spinal alignment. Our characterization of Ebf2 knock-out mice revealed a requirement for Ebf2 in the differentiation of a subset of MMC MNs, indicating molecular diversity within MMC neurons. Intriguingly, transgenic expression of mEbf1 or mEbf2 can rescue axial MN differentiation and locomotory defects in nematodes (*Caenorhabditis elegans*) lacking *unc-3*, the sole *C. elegans* ortholog of the COE family, suggesting functional conservation among mEbf1, mEbf2 and nematode UNC-3.

**Conclusions:** These findings support the hypothesis that the genetic programs controlling axial MN development are deeply conserved across species, and further advance our understanding of such programs by revealing an essential role for Ebf2 in mouse axial MNs. Because human mutations in COE ortholgs lead to neurodevelopmental disorders characterized by motor developmental delay, our findings may advance our understanding of these human conditions.

## BACKGROUND

The mammalian neuromuscular system is essential for distinct motor behaviors ranging from locomotion and dexterity to basic motor functions, such as breathing and maintenance of spinal alignment [1]. The underlying basis for achieving these diverse outputs lies in the assembly of distinct neuronal circuits dedicated to control different muscles. In the mouse spinal cord, for example, these circuits are composed of various motor neuron (MN) subtypes organized into distinct clusters of cells (termed “columns”) along the rostrocaudal axis (**Fig. 1A**). At the brachial and lumbar levels, MNs of the lateral motor column (LMC) innervate limb muscles, which are essential for locomotion and dexterity [2, 3]. Breathing is controlled by cervical MNs of the phrenic motor column (PMC) that innervate the diaphragm, and by thoracic MNs of the hypaxial motor column (HMC) that innervate hypaxial (intercostal and abdominal) muscles. In contrast to these segmentally-restricted columns (LMC, PMC, HMC), MNs of the medial motor column (MMC) are generated along the entire length of the spinal cord and innervate epaxial (back) muscles necessary for maintenance of spinal alignment [1]. (**Fig. 1A, C**). While remarkable progress has been made in deciphering the molecular mechanisms that specify limb-innervating MNs (LMC), the genetic programs underlying the development of hypaxial (HMC) and epaxial (MMC) muscle-innervating MNs are poorly studied [1].

**Figure 1:**
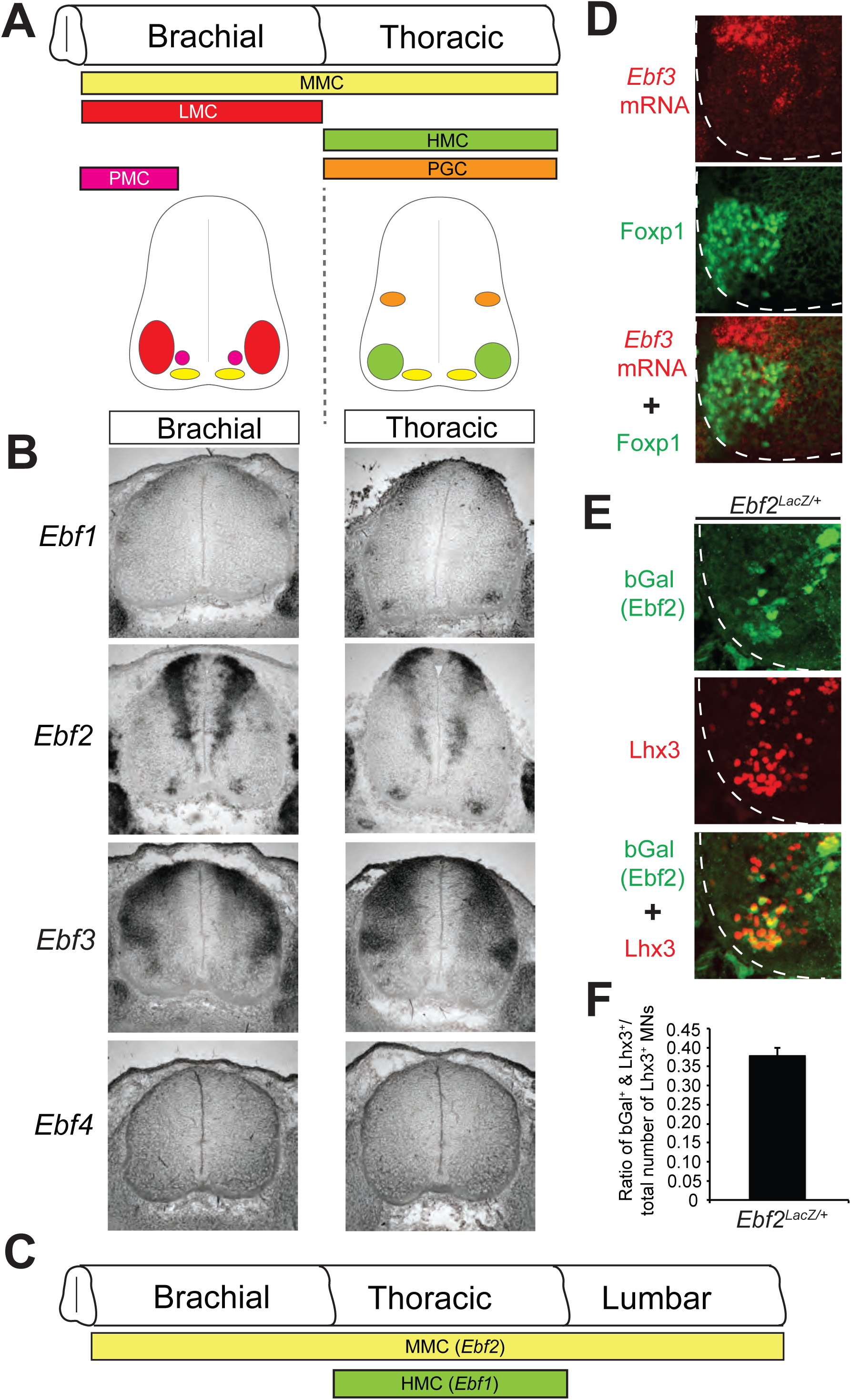
Ebf1 and Ebf2 are expressed in axial muscle-innervating motor neurons. **A**: Schematic of the spinal cord showing different columns of MNs (color-coded) at distinct regions along the rostrocaudal axis (brachial and thoracic). A cross-section of each region is provided below. **B**: RNA ISH analysis for mouse Ebf1-Ebf4 at e13.5 of WT spinal cords. **C**: Schematic summarizing the expression of *Ebf1* (HMC) and *Ebf2* (MMC) based on data from panel B. **D**: Antibody staining for the LMC marker (Foxp1, green signal) combined with fluorescent RNA ISH for *Ebf3* (red signal) revealed no co-localization in WT e13.5 spinal cord. N=3. **E-F**: Double immunostaining for Lhx3 (MMC marker in red) and bGal (*Ebf2* reporter in green) shows that ~40% of MMC neurons express *Ebf2*. Quantification (**F**) provided as the ratio of MNs that are double positive for Lhx3 and bGal over the total number of Lhx3 positive MNs. For this analysis, *Ebf2 ^LacZ/+^* embryos were used at e12.5. N=5.

The identity of MN subtypes that belong to a specific column, termed “columnar identity”, is defined by combinatorial expression of transiently expressed TFs [2, 4, 5]. For example, LMC columnar identity is imposed by co-expression of several LIM homeodomain proteins (Islet1, Islet2, Hb9 [Mnx1]) and the Forkhead box protein P1 (Foxp1). On the other hand, combinatorial expression of Islet1, Islet2, Lhx3 and Hb9 demarcates MMC columnar identity [6]. The highly conserved Hox proteins also play key roles in establishing columnar MN identity. Distinct combinations of Hox genes are expressed in MNs along the rostrocaudal axis of the spinal cord [3, 7]. At brachial and lumbar levels, for example, Hox genes control LMC identity through the direct activation of Foxp1 [48, 49]. However, the identity of the non-segmentally restricted MMC neurons appears to be insensitive to the activities of Hox proteins [1, 8], suggesting that distinct genetic programs have evolved for the establishment of limb-innervating (LMC) and axial muscle-innervating (MMC) MNs.

A large body of work on limb-innervating MNs (LMC) demonstrated that combinatorial TF expression is also necessary for further diversification of MN subtypes within the same column, providing evidence for molecular diversity within the LMC column [4, 5, 9]. In the mouse hind limb, co-expression of Islet1, Islet2, Er81 and Nkx6.1 is necessary for specification of MNs located in medial LMC (LMCm) that innervate ventral limb muscles. On the other hand, co-expression of Hb9, Lhx1, Islet2, Er81 and Pea3 defines lateral LMC (LMCl) identity, ensuring proper innervation of dorsal limb muscles. Unlike LMC MNs, it remains unclear to date whether molecular diversity exists within the axial muscle-innervating MNs. While combinatorial expression of Islet1, Islet2, Lhx3 and Hb9 demarcates MMC columnar identity [1, 6], molecular markers that label subsets of MMC neurons have not been identified in mice, thereby limiting our ability to genetically access and study these neurons. In other words, is the anatomical diversity found within MMC MN subtypes accompanied by molecular diversity? Similar to the MMC case, the genetic programs underlying the differentiation of hypaxial muscle-innervating MNs (HMC) are poorly studied, in part due to the lack of HMC-specific molecular markers.

MMC neurons are thought to represent the ancestral “ground state” from which all other spinal cord MN subtypes subsequently evolved [2]. This notion is supported by the fact that “MMC-like” neurons innervate axial/body-wall muscles in limbless vertebrates (e.g., lamprey), insect larvae (e.g., *Drosophila melanogaster*) and nematodes (e.g., *Caenorhabditis elegans*), suggesting that an MMC-like population likely represents the ancestral condition of MNs in bilaterians [1]. It is therefore not surprising that at least some of the intrinsic genetic programs of axial MN development appear to be conserved from invertebrates to vertebrates. Supporting this notion, “MMC-like” neurons in *Drosophila* and *C. elegans* can be defined, similar to vertebrates, by the expression of Hb9, Lhx3 and Islet1/2 orthologs [10]. Cross-species comparisons have also revealed “lost” mechanisms. This is perhaps best exemplified by the case of the conserved homeodomain TF *even-skipped (eve)*. *Drosophila eve* and its *C. elegans* ortholog *vab-7* are utilized for specification of body wall muscle-innervating MNs [11-15], while the *eve/vab-7* mouse ortholog Evx1 is not involved in MN specification. Instead, Evx1 is strictly required for V0 spinal interneuron fate [16]. Although axial MNs are used for distinct motor functions in different species (locomotion in limbless vertebrates, insect larvae and nematodes *versus* maintenance of spinal alignment in mammals), the aforementioned examples collectively illustrate that cross-species comparisons can reveal the extent of conservation in the genetic programs underlying axial MN development.

Our previous studies in the nematode *C. elegans* revealed that UNC-3, the sole ortholog of the Collier/Olf/Ebf (COE) family of TFs, is required for differentiation of body wall muscle-innervating MNs that control locomotion [17-19]. COE family orthologs are expressed in the nervous system of very distant species ranging from cnidarians (e.g., sea anemone) [20] to bilaterians (nematodes [21-23], flies [24], frogs [25], zebrafish [26], mice [27-33]), indicating an ancient role in nervous system development. Functional studies have shown that the sole *Drosophila* COE ortholog *collier (or knot)* is required for peptidergic neuron specification [34-37], and COE orthologs in frog and chick embryos function to promote neuronal differentiation [25, 38]. Four COE orthologs are embedded in the mouse genome, mEbf1-mEbf4. Previous reports have identified mEbf1 as a key player in facial MN migration, as well as neuronal differentiation in the striatum and retina [29, 30, 39]. Mouse Ebf2 is required for neuronal cell migration and differentiation in the cerebellum and olfactory epithelium, where it is believed to function in a partially redundant manner with Ebf3 [27, 28, 33, 40, 41]. However, the function of mouse Ebfs in spinal MN development remains poorly understood.

Our previous studies on cholinergic MNs of *C. elegans* and the simple chordate *Ciona intestinalis* indicated that the function of UNC-3 is conserved from nematodes to simple chordates [19]. Here, we provide evidence that the function of UNC-3 may be conserved from *C. elegans* to mammals. We found, in mice, that mEbf1 and mEbf2 are implicated in axial MN development. In the embryonic mouse spinal cord, mEbf1 is selectively expressed in hypaxial muscle-innervating MNs (HMC), while mEbf2 is expressed in epaxial muscle-innervating MNs (MMC). Using Ebf2 KO mice, we assessed in vivo the function of Ebf2 and revealed its requirement for differentiation of a subset of MMC neurons. Lastly, cross-species transgenic rescue experiments demonstrated that mouse Ebf1 or Ebf2 can functionally substitute for nematode UNC-3. Altogether, our study uncovers an ancient role for COE family TFs in axial MN development. Because human mutations in COE orthologs cause neurodevelopmental disorders characterized by motor developmental delay [17, 42-46], our findings could help advance our understanding of these conditions.

## METHODS

### Mouse husbandry

All mouse procedures were approved by the Institutional Animal Care and Use Committee (IACUC) of the University of Chicago.

### *RNA in situ hybridization,* antibody staining and quantification

Embryos were harvested at e12.5, e13.5, and e15.5, fixed in 4% paraformaldehyde for 1.5–2 h, placed in 30% sucrose over-night (4°C), and embedded in optimal cutting temperature (OCT) compound. Cryosections were generated and processed for in situ hybridization or immunohistochemistry using the antibodies against Foxp1, Lhx3, Hb9, and bGal as previously described[9, 47]. Images were obtained with a high-power fluorescent microscope (Zeiss Imager.V2) and analyzed with Fiji software[48]. Cell counting was performed using ImageJ software in at least three embryos per genotype. Cells stained for Foxp1, Lhx3, Hb9, bGal (Ebf2) were counted at the same position along the spinal cord for each genotype using the Cell count plug-in. The axial projections of Hb9-GFP labeled MNs were visualized in vibratome sections (150µm) of control (*Hb9-GFP*) and *Ebf2 KO; Hb9-GFP* spinal cords at e12.5.

### C. elegans strains

Nematode *C. elegans* strains were maintained as previously described[49]. Two strong loss-of-function (putative null) alleles (*e151, n3435*) for *unc-3* were used in this study[22]. Cholinergic MN reporters used: *juIs14 [acr-2^prom^::gfp], vsIs48 [unc-17^prom^::gfp]*. The following rescuing transgenic strains were generated: two rescuing transgenic lines for *unc-3^prom^::mEbf1::unc-54 3’UTR* and two rescuing transgenic lines for *unc-3^prom^::mEbf2::unc-54 3’UTR*.

### Rescue experiments in *C. elegans*

The *unc-3^prom^::mEbf1::unc-54 3’UTR* construct and the *unc-3^prom^::mEbf2::unc-54 3’UTR* construct were generated to contain a 558bp proximal fragment (immediately upstream of ATG) of the *unc-3* promoter amplified from *C. elegans* wild type (N2) genomic DNA. In addition, each construct contains corresponding *Ebf1* or *Ebf2* coding sequences amplified from mouse cDNA and fused to *C. elegans unc-54 3’UTR*. Fragments were assembled through the Gibson Assembly method, which can be found at https://www.neb.com/applications/cloning-and-synthetic-biology/dna-assembly-and-cloning/gibson-assembly. These constructs were injected at 25 ng/µl into both *unc-3(n3435); juIs14 [acr-2^prom^::gfp] and unc-3(e151); vsIs48 [unc-17^prom^::gfp]* mutant strains together with *myo-2^prom^::GFP* (co-injection marker) and a carrier plasmid that serves as filler DNA (pBluescript). At least two stable transgenic lines were established for each construct and tested for rescue of the *unc-3* mutant phenotypes.

### Body bend/thrashing assay

Young-adult nematodes (2 days old) were transferred to an NGM plate containing a droplet (100ul) of M9 buffer. After 1 min of adaptation, the number of body bends for 30 sec was quantified as described in[50]. *A movement of the worm that swings its head and/or tail to the same side is counted as one thrash.*

### Phylogenetic tree for COE family members

The peptide sequence for UNC-3 and Collier (Knot) were recovered from WormBase and FlyBase, respectively. The peptide sequences for mEbf1, mEbf2, mEbf3, mEbf4, and Ciona intestinalis COE were downloaded from Ensembl Genome Browser. While a single isoform was found for UNC-3, Collier and Ciona COE, several protein isoforms for mouse Ebfs were reported in Ensembl. The longest mEbf1, mEbf2, mEbf3, and mEbf4 isoforms were considered for the generation of the phylogenetic tree shown in Figure 4A. Phylogenetic analysis was performed at http://phylogeny.lirmm.fr using the “one-click” mode [51-53].

### Statistical analysis

For data quantification, graphs show values expressed as mean ± standard deviation (SD). Statistical significance was determined with the unpaired two-tailed Student’s t test using Microsoft Excel.

## RESULTS

### Ebf1 and Ebf2 are expressed in distinct MN subtypes that innervate axial muscles

Since orthologs of COE family TFs control aspects of MN differentiation in the nematode *C. elegans* and the simple chordate *C. intestinalis* [19], we hypothesized that the function of COE TFs is conserved in mouse MNs. Four orthologs of the COE family are embedded in the mouse genome (mEbf1 - mEbf4). We therefore sought to examine the expression pattern of all four mEbfs in the developing mouse spinal cord through RNA in situ hybridization (ISH). Since different columns (MMC, HMC, LMC, PMC, PGC) are found along the rostro-caudal axis of the spinal cord, we performed RNA ISH for Ebf1-4 at brachial, thoracic, and lumbar levels (**Fig. 1A**). We chose embryonic day 13 (e13.5) as the most appropriate time point for this analysis because post-mitotic MNs by e13.5 are organized into columns that can be distinguished with molecular markers (**Fig. 1A**). We found that mEbf1 is expressed in MNs of the HMC but no other column at brachial, thoracic, or lumbar levels, constituting a highly specific, post-mitotic marker for HMC neurons (**Fig. 1B**). Unlike mEbf1, mEbf2 is selectively expressed in MNs along the entire length of the spinal cord that belong to MMC (**Fig. 1B, C**). Our RNA ISH revealed expression of mEbf3 in the ventral zone of e13.5 spinal cords, a territory closed to the location of LMC MNs. However, coupling of mEbf3 RNA ISH analysis with antibody staining for Foxp1, a LMC-specific marker, revealed no co-localization (**Fig. 1D**). We detected though mEbf3 expression in interneurons of the intermediate and dorsal zones, as well as dorsal root ganglia (DRG) (**Fig. 1B, Suppl. Fig. 1**). We note that mEbf1 and mEbf2 expression was also detected in dorsal interneurons and DRG neurons (**Fig. 1B, Suppl. Fig. 1**). No expression for mEbf4 was detected in spinal cord and DRG neurons at e13.5 (**Fig. 1B, Suppl. Fig. 1**). These findings extend previous observations of the Ebf1-3 expression pattern in the spinal cord performed at earlier stages, before MNs are organized into columns [30]. By examining the expression pattern of all members of the COE family of TFs at e13.5 spinal cords, we demonstrate that mEbf1 and mEbf2 are selectively expressed in axial muscle-innervating MNs of the HMC and MMC, respectively. Based on molecular and positional criteria, we found no evidence of Ebf expression at e13.5 in the remaining motor columns, such as the limb-innervating MNs of the LMC.

### mEbf2 is required for the differentiation of a subset of MMC neurons

The mEbf2 expression in MNs along the spinal cord (MMC) is reminiscent of the UNC-3 expression pattern in body wall muscle-innervating MNs along the *C. elegans* nerve cord [19, 23]. We therefore hypothesized that – similar to its *C. elegans* ortholog (UNC-3) – mEbf2 is also required for differentiation of mouse MNs (MMC) that innervate body wall muscles, and more precisely epaxial muscles. Before testing this hypothesis, it is necessary to determine whether all MMC neurons or a fraction of them express mEbf2 (**Fig. 1B**). To this end, we used a mouse line, which carries a promoterless lacZ cDNA cassette to replace the first five exons of the endogenous Ebf2 locus [40]. Heterozygous Ebf2 ^LacZ/+^ mice enabled us to perform double immunofluorescence staining for b-galactosidase (produced by Lacz), which serves as an Ebf2 reporter, and the MMC-specific marker Lhx3 [6]. Following quantification of the number of MNs that co-express b-galactosidase (Ebf2) and Lhx3 at e13.5, we found that ~ 40% of the MMC neurons express Ebf2 at any given region (brachial, thoracic, or lumbar) of the spinal cord (**Fig. 1E-F**). This finding corroborates our RNA ISH analysis for mEbf2 expression in MMC (**Fig. 1B**), and further suggests that molecular differences do exist within MMC neurons.

To examine the function of Ebf2, we used Ebf2 ^LacZ/LacZ^ homozygous mice. These mice globally inactivate Ebf2 gene activity [40], and will be referred to as Ebf2 KO mice hereafter. Since orthologs of Ebf2, in nematodes and simple chordates, are required for expression of the highly conserved genes involved in acetylcholine (ACh) biosynthesis, we evaluated the expression of the ACh pathway genes VAChT and Acly by RNA ISH in Ebf2 KO embryonic spinal cords (**Fig. 2A**). We did not detect any differences, a finding that raises the possibility of genetic redundancy among the 4 mEbf factors. However, RNA ISH analysis did not show any expression of Ebf1, Ebf3, and Ebf4 in MMC neurons of Ebf2 KO embryos (**Fig. 2B)**. We therefore conclude that – unlike its nematode and simple chordate orthologs – mEbf2 does not disrupt the expression ACh pathway genes.

**Figure 2:**
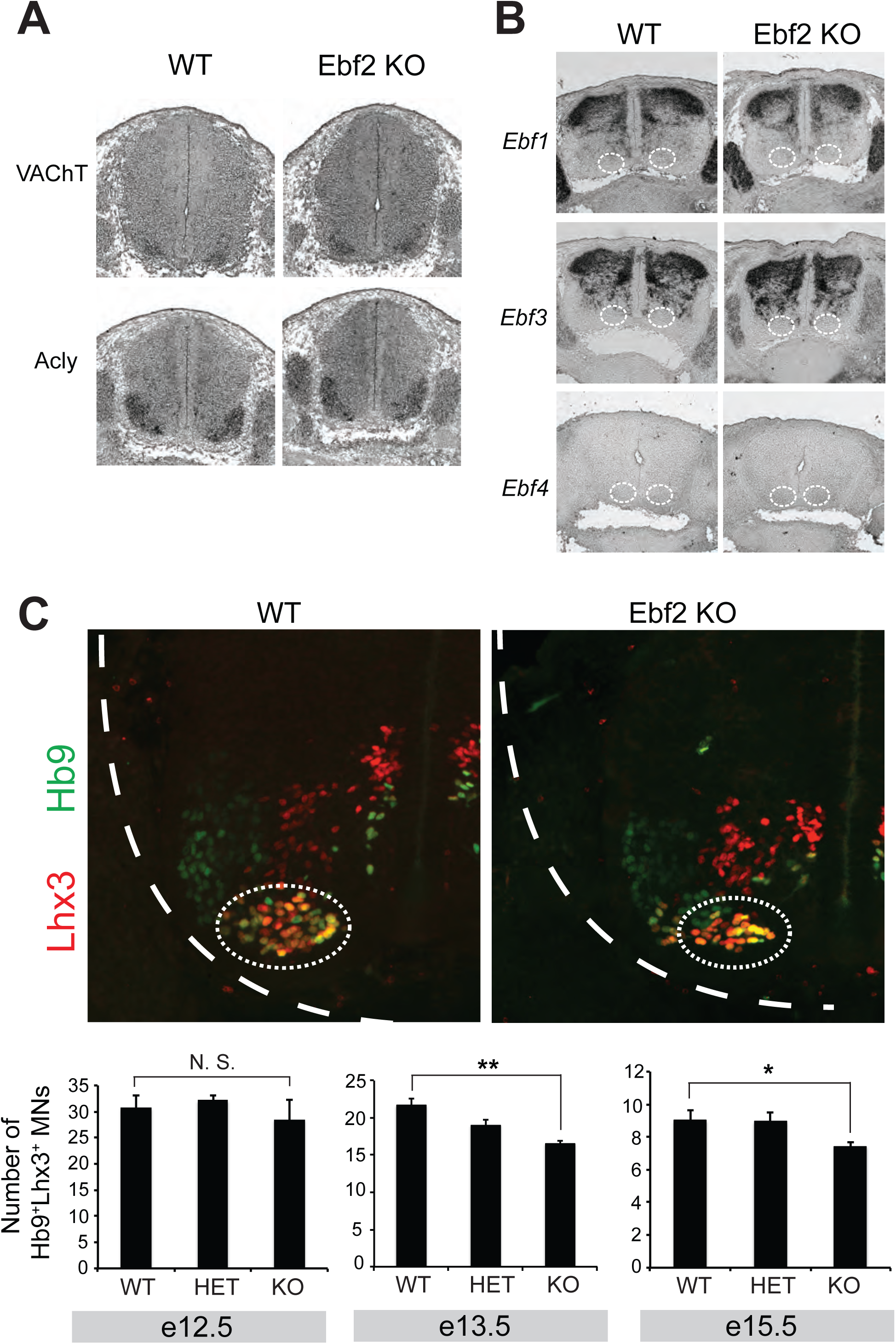
Characterization of MMC neurons in Ebf2 KO animals. **A**: The expression of genes involved in ACh biosynthesis (VAChT, Acly) is not affected in MMC neurons of Ebf2-KO spinal cords at e13.5. N=3. **B**: RNA ISH analysis on Ebf2-KO spinal cords at e12.5 or e13.5 reveals no compensatory upregulation of *Ebf1, Ebf3* and *Ebf4* transcripts in MMC neurons. N=3. **C**: The number of MMC neurons co-labeled with Lhx3 (red) and Hb9 (green) is not affected in Ebf2 KO spinal cords at e12.5, but is significantly decreased at e13.5 and e15.5. Student’s t-test was performed. *: p < 0.05, **: p < 0.01. N=4-7. In B and C, the location of MMC neurons is indicated with a dotted white circle.

The expression of genes involved in ACh biosynthesis is a common feature shared by all spinal MNs irrespective of columnar identity. We then asked whether molecular features specific to MNs of the MMC column are affected in Ebf2 KO embryos. It is well established that co-expression of the TFs Lhx3 and Hb9 demarcates the MMC neurons in the developing spinal cord [6, 54]. We quantified the number of double-positive (Lhx3+ and Hb9+) MNs in the ventral zone of WT, Ebf2 heterozygous (Ebf2 ^LacZ/+^) and homozygous KO (Ebf2 ^LacZ/LacZ^) spinal cords at e12.5, e13.5, and e15.5 (**Fig. 2C**). Although no differences were observed at e12.5, we found a statistically significant decrease in the number of MMC MNs (Lhx3+ and HB9+) in both Ebf2 KO spinal cords at e13.5 and e15.5, suggesting that Ebf2 is required for later stages of MMC neuron differentiation (**Fig. 2C**). This possibility is further supported by the fact that Ebf2 expression in MMC is maintained throughout embryonic development (Ebf2 ISH at e18, **Suppl. Fig. 1**). Next, we examined the expression of two recently described molecular markers for MMC fate [8] (Dkk3, Ldb2). We found no differences when compared their expression in WT and Ebf2 KO spinal cord at e13.5. Lastly, we asked whether these molecular defects are accompanied by anatomical defects in MMC neurons, such as the ability to reach their epaxial muscle targets. To this end, we visualized all spinal MN axons (including the MMC axons) using the Hb9-GFP mouse line [55], which we crossed to Ebf2 KO mice. We were unable to detect any differences in the morphology or thickness of the MMC axonal bundle (nerve) that reaches epaxial muscles (**Fig. 3A**). We note, however, that the Hb9-GFP mouse line labels with GFP all MMC neurons, including the 40% of them that express Ebf2. This lack of specificity could contribute to our inability to detect any putative MMC axonal defects in Ebf2 KO mice. Nevertheless, the above molecular analysis suggests that Ebf2 is required for differentiation of a subset of MMC neurons.

**Figure 3.**
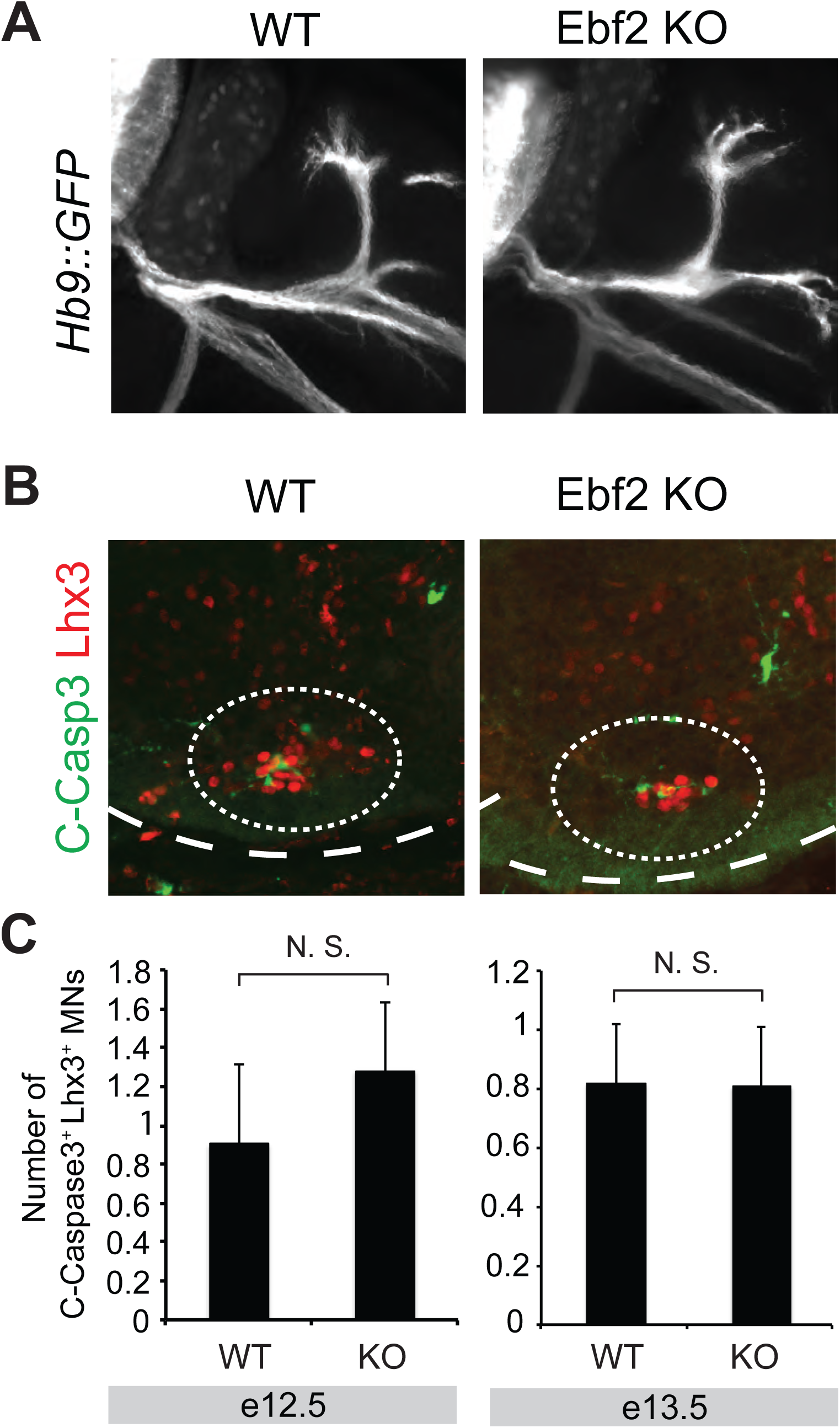
No evidence for axon guidance defects and Caspase 3-mediated cell death in MMC neurons of Ebf2 KO mice. **A**: Motor neuron axons (bundled into nerves) are visualized with the Hb9::GFP transgene in WT and Ebf2 KO spinal cords at e12.5. The MN axons that belong to MMC are shown with an arrow. **B**: Representative images of WT and Ebf2 KO spinal cords (cross-sections) immunostained with antibodies against Cleaved-Caspase 3 (C-Casp3 in green) and Lhx3 (MMC marker in red) at e12.5. The location of MMC neurons is circled. **C**: Quantification of the number of MMC neurons (Lhx3^+^) that are also positive for C-Casp3 staining in WT and Ebf2 KO spinal cords. This analysis was performed at e12.5 and e13.5. N. S. Not Significant differences were found when Student’s t-test was performed. N = 4.

### The MMC differentiation defect in Ebf2 KO animals is not a result of Caspase 3-mediated cell death

The reduction in the number of MMC neurons co-expressing the molecular markers Lhx3 and Hb9 in Ebf2 KO embryos could be attributed either to a *bona fide* differentiation defect (i.e., MMC neurons are normally generated in the absence of Ebf2 gene activity, but fail to express MMC-specific markers), or to a developmental defect that affects the number of MMC neurons (e.g., loss of Ebf2 triggers apoptosis in MMC neurons, thereby reducing their number). To investigate the latter possibility, we quantified the number of MMC neurons that express cleaved caspase-3 (C-Casp3), a well-documented pro-apoptotic marker and the converging point of several pro-apoptotic signaling pathways [56]. Upon double staining for Lhx3 (MMC marker) and C-Casp3, our quantification analysis at e12.5 and e13.5 revealed no statistically significant differences between WT (Ebf2 ^+/+^) and Ebf2 KO embryos (**Fig. 3B-C**). Although there is a possibility that we missed a narrow time window (between e12.5 and e13.5) during which increased apoptosis occurs, our C-Casp3 data thus far suggest that the MMC differentiation defect in Ebf2 mutant mice is not due to C-Casp3-dependent cell death that could affect the generation of MMC neurons. We note that this is also the case for nematodes lacking the Ebf2 homolog *unc-3*; axial MNs are normally generated in these mutant animals but fail to express MN-specific differentiation markers [19].

### Mouse Ebf1 and Ebf2 rescue MN differentiation defects in *unc-3* mutant nematodes

Our previous studies in the nematode *C. elegans* and the chordate *Ciona intestinalis* suggested that the function of UNC-3 in controlling axial MN differentiation is conserved from nematodes to simple chordates [19]. Here, we investigated whether the function of UNC-3 is conserved from nematodes to mammals through cross-species rescue experiments. Given that mEbf1 and mEbf2 are expressed in mouse axial MNs (**Fig. 1A-C**) and are phylogenetically closer to UNC-3 (based on sequence similarity) than mEbf3 and mEbf4 (**Fig. 4A)**, we sought to explore the functional conservation of mEbf1 and mEbf2 and nematode UNC-3. To this end, we expressed the mEbf1 or mEbf2 coding sequence specifically in MNs of *unc-3* mutant nematodes that carry strong loss-of-function or null *unc-3* alleles (*e151, n3435*) [22]. As assessed by analyzing the expression of two *unc-3*-dependent MN differentiation markers (*acr-2: <u>AC</u>h <u>r</u>eceptor subunit 2; unc-17: <u>v</u>esicular <u>ACh t</u>ransporter [VAChT])[19],* we found that both mEbf1 and mEbf2 could rescue the *unc-3* mutant phenotype and restore *acr-2* and *unc-17* expression (**Fig. 4B-C**). In addition to evaluating expression of MN-specific molecular markers, we asked whether MN-specific expression of mEbf1 or mEbf2 can rescue the severe locomotory defects previously observed in *unc-3* null nematodes [49]. To test this possibility, we employed a well-established thrashing assay[50] and counted the number of body bends per 30 sec when adult worms are placed into solution (M9 buffer) (**Fig. 4D**). Our quantification revealed that MN-specific expression of either mEbf1 or mEbf2 in *unc-3* null animals resulted in a statistically significant increase in the number of body bends/30sec when compared to *unc-3* null animals that do not carry the mEbf1 or mEbf2 rescuing transgenes (**Fig. 4D**). Apart from corroborating the cell autonomy of the *unc-3* mutant phenotype (because mEbf1 and mEbf2 were expressed under the control of a MN-specific promoter [*unc-3 ^558bp prom^*]), our behavioral and molecular analysis (*acr-2, unc-17*) show that mEbf1 and mEbf2 can functionally compensate for the lack of nematode *unc-3*.

**Figure 4.**
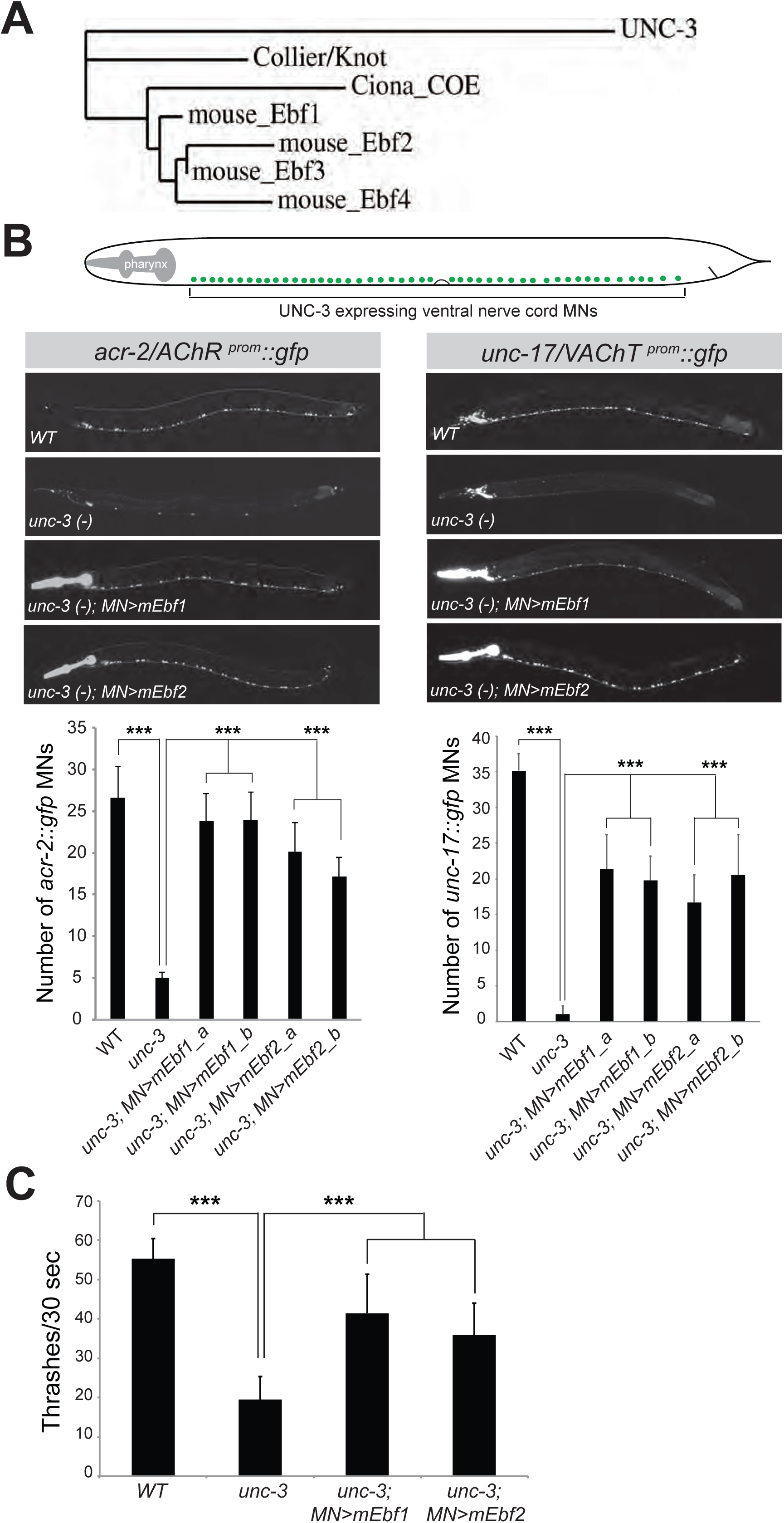
Mouse Ebf1 and Ebf2 rescue MN specification and locomotory defects in nematodes lacking *unc-3*. **A**: Phylogenetic tree generated using Phylogeny.fr [51-53]. **B**: Schematic showing the cell body position of ventral nerve cord MNs (labeled with green) in *C. elegans.* Forty (40) of these MNs are cholinergic and express UNC-3. The expression of two cholinergic MN markers (*acr-2*/AChR, *unc-17*/VAChT) is dramatically reduced in *unc-3* mutant animals carrying loss-of-function alleles (*e151, n3435*). Transgenic expression of mEbf1 *[Punc-3::mEbf1::unc-54 3’ UTR]* or mEbf2 *[Punc-3::mEbf2::unc-54 3’ UTR]* significantly restores the MN expression of *acr-2*/AChR and *unc-17*/VAChT. Two transgenic lines per construct were used. A representative image of one rescuing line is shown. Fifteen animals at larval stage 4 (L4) were imaged for each genotype. Note that the fragment of *unc-3* promoter (558bp) used for rescue drives expression of mEbf1 or Ebf2 in all *acr-2* positive MNs (~35 MNs), resulting in significant rescue. However, this *unc-3* promoter fragment (558bp) drives expression of mEbf1 or Ebf2 in ~30 out of the 40 MNs that express *unc-17* in the ventral nerve cord, thereby resulting in partial rescue. **C**: Quantification of the results shown in panel B. Two transgenic lines (indicated with a and b) per rescuing construct were quantified. Student’s t-test was performed. N=15. *: p < 0.05, **: p < 0.01, ***: p < 0.001. **D**: Thrashing assay shows that the movement defects of *unc-3* mutant nematodes can be significantly restored upon MN-specific expression of mEbf1 or mEbf2. Two day-old adult nematodes were used. N=10. Student’s t-test was performed. *: p < 0.05, **: p < 0.01, ***: p < 0.001.

## DISCUSSION

While there is a remarkable wealth of studies focusing on molecular programs controlling the differentiation of limb-muscle innervating MNs, the development of axial muscle-innervating MNs has received much less attention [1]. In this paper, we provide evidence for an ancient role of the COE family of TFs in axial MN development. First, we find that mEbf1 is a novel post-mitotic marker for HMC neurons, which innervate hypaxial muscles necessary for breathing, opening the door for future studies aiming to illuminate the poorly explored genetic programs of HMC specification. Second, this study advances our understanding of the genetic programs underlying axial MN development: (**i**) By uncovering an essential role for mEbf2 in the differentiation of MMC neurons, which innervate epaxial muscle and control spinal alignment, and (**ii**) By revealing that molecular diversity does exist within MMC neurons as Ebf2 is expressed in a significant fraction (~40%) of these MNs. Third, our cross-species transgenic rescue experiments provide evidence for functional equivalence between COE orthologs, suggesting that the function of these TFs in the development of axial muscle-innervating MNs is conserved from nematodes to mammals. Altogether, this study indicates that a viable approach for advancing our understanding of axial MN development may lie in comparative studies of the genetic programs that specify invertebrate and vertebrate MNs.

Comparison of the role of COE family TFs in MNs of nematodes, simple chordates, and mice (this study) suggests that COE TFs played an ancient role in axial muscle-innervating MNs. In the nematode *C. elegans*, the sole COE ortholog UNC-3 is expressed in post-mitotic MNs that innervate axial muscle [19, 23]. This is also the case for the sole COE ortholog in the simple chordate, *Ciona intestinalis* [19]. Interestingly, the zebrafish COE ortholog *Zcoe2* appears to also be expressed in axial muscle-innervating MNs [26]. Here, we describe that mEbf1 and mEbf2 are respectively expressed in HMC and MMC post-mitotic mouse MNs, which innervate distinct groups of axial muscle (HMC neurons > hypaxial muscle; MMC neurons > epaxial muscle). It has been suggested that HMC and MMC MNs likely reflect the vestige of an ancestral spinal motor column organization from which other motor columns derived in vertebrates [1, 2]. During this process, we propose that COE ortholog expression was maintained in axial muscle-innervating MNs (HMC and MMC), but lost in other motor columns, such as limb-innervating MNs (LMC). A speculative interpretation of our findings is that COE-type TFs may have controlled the specification of axial muscle-innervating MNs in the common ancestor of nematodes, simple chordates and vertebrates. It has been proposed that the ancestor to all bilaterians had a fairly complex nervous system and likely used an axial-like locomotor circuit to move [1, 57, 58].

The function of axial muscle-innervating MNs is different between vertebrates and invertebrates [1]. For example, axial muscle-innervating MNs in *C. elegans* are required for locomotion, whereas, in mice, HMC and MMC neurons have adopted a more specialized role (HMC, breathing; MMC, maintenance of spinal alignment). To accommodate this functional change, it is conceivable that during evolution modifications have occurred in the molecular programs that specify axial muscle-innervating MNs. At least two, non-mutually exclusive scenarios can be envisioned: (**i**) New, vertebrate-specific programs may have evolved for the control of axial MN differentiation in vertebrates, or (**ii**) Modifications in pre-existing genetic programs found in invertebrate axial MNs may have occurred to accommodate the more specialized role of axial muscle-innervating MNs in vertebrates. Our findings lend support to both of these scenarios. In *C. elegans* axial MNs, UNC-3 is required for axon guidance, ACh biosynthesis, expression of conserved MN-specific TFs (e.g., *ceh-12*/Hb9 or Mnx1), and induction of a large battery of MN identity-defining genes (e.g., ion channels, neuropeptides, NT receptors) [19]. However, our analysis of mice lacking Ebf2 (UNC-3 ortholog) revealed a more specialized role for Ebf2 in axial muscle-innervating MNs. We did not observe axon guidance defects or reduction in expression of genes critical for ACh biosynthesis (e.g., VAChT, Acly) in MMC neurons of Ebf2 KO mice. Although Ebf2 is able to induce VAChT gene expression in *C. elegans* MNs (**Fig. 4**), this ability has been lost in mouse MMC neurons (**Fig. 2A**), which appear to have evolved a new genetic program orchestrated by Islet1 for the control of ACh biosynthesis genes [59]. However, we found that – similar to *unc-3* mutant nematodes – the number of axial MNs expressing the conserved MN determinant Hb9 is significantly reduced in Ebf2 KO mice, indicating that aspects of the pre-existing, UNC-3-mediated genetic program have been conserved.

A current limitation in our Ebf2 KO analysis is the lack of additional molecular markers for MMC neuronal identity (e.g., ion channels, neuropeptides, NT receptors), which prevents the characterization of Ebf2 function in a comprehensive manner. As such, future molecular profiling studies are needed to generate MMC-specific markers that will enable us to determine the degree of functional conservation between mouse Ebf2 and nematode UNC-3 in axial MN development. A potential outcome of such profiling studies is the identification of genes that are either expressed in all MMC neurons or in specific subsets of them, as suggested by our observation that ~40% of MMC neurons express Ebf2. The availability of such markers would allow the study of the extent of molecular diversity within the MMC neurons, which is presently unknown [1].

Our comparative study has evo-devo implications pertaining to the long-standing question of whether conservation of TF expression in orthologous cell types of different species is accompanied by conservation of TF function in these cell types. Through cross-species transgenic rescue experiments, we found that mEbf1 and mEbf2 can functionally substitute for *unc-3* function in *C. elegans* MNs, as assessed by monitoring expression of two MN identity markers (*acr-2*/AChR and *unc-17*/VAChT). Quite remarkably, mEbf1 and mEbf2 were also able to rescue the severe locomotory defects observed in *unc-3* nematodes, indicating that most, if not all, UNC-3 target genes are normally expressed. These findings support the view that during evolution the structure and domain architecture of COE TFs did not significantly diverge. Indeed, there is high similarity between the DNA-binding and helix-loop-helix (HLH) domains of UNC-3, mEbf1 and mEbf2 [20, 26]. Moreover, a phylogenetic comparison among all 4 mouse Ebfs and UNC-3 revealed that mEbf1 and mEbf2 are more closely related to UNC-3 (**Fig. 4A**). We therefore surmise that the different functions of nematode UNC-3 and mouse Ebf2 in axial muscle-innervating MNs, i.e., UNC-3, but not Ebf2, controls ACh biosynthesis, may have arisen due to divergence at the level of their target genes (e.g., loss of unique sets of UNC-3 target genes).

## CONCLUSIONS

Neuronal control of muscle associated with the central body axis is an ancient and essential function of both invertebrate and vertebrate nervous systems. Here, we provide evidence for an ancient role of the COE family of TFs in axial MN development. In the future, it will be interesting to examine whether other evolutionarily conserved TFs (e.g., *bnc-1*/BNC, *cfi-1*/Arid3a) that work together with UNC-3 to determine axial MN differentiation in *C. elegans* also control aspects of axial MN development in mice [17]. Further cross-species comparisons of the genetic programs underlying axial MN development may provide valuable insights into how axial MN diversity is generated and distinct functions of axial MNs have evolved across the animal kingdom. Beyond MNs, our study highlights the usefulness of cross-species comparisons for the identification of genetic programs that orchestrate neuronal fate specification.

## List of abbreviations

MN4: Motor neuron
TF: Transcription factor
LMC: Lateral motor column
HMC: Hypaxial motor column
MMC: Medial motor column
WT: wild type
KO mice: Knock-out mice

## Acknowledgements

Some *C. elegans* strains were provided by the *Caenorhabditis* Genetics Center (CGC), which is funded by NIH Office of Research infrastructure Programs (P40 OD010440). The antibodies for Foxp1, Lhx3, and Hb9 were a gift from the lab of Thomas Jessell. We thank Ellie Heckscher, Robert Carrillo and members of the Kratsios lab for providing comments on the manuscript. This work was supported by an NINDS grant (R00NS084988) and a Whitehall Foundation grant to P.K.

---

**Suppl. Fig 1: Additional characterization of Ebf expression in the mouse spinal cord. A**: RNA ISH at e13.5 reveals Ebf1, Ebf2 and Ebf3 expression in DRG neurons (red circles). **B**: Double immunofluorescence staining for bGal (Ebf2 reporter in green) and Lhx3 (MMC marker in red) reveals co-localization (arrows) at a late embryonic stage (e18.5). A representative image is shown from the brachial region. A subset of Lhx3 positive neurons express bGal (Ebf2), which is also the case at earlier (e13.5) stages (shown in Figure 1).

